# High prevalence of S. pyogenes Cas9-specific T cell sensitization within the adult human population – A balanced effector/regulatory T cell response

**DOI:** 10.1101/295139

**Authors:** Dimitrios L. Wagner, Leila Amini, Desiree J. Wendering, Petra Reinke, Hans-Dieter Volk, Michael Schmueck-Henneresse

**Affiliations:** Institute for Medical Immunology, Charité - University Medicine Berlin, Germany; Department of Nephrology and Internal Intensive Care, Renal and Transplant Research Unit, Charité - University Medicine Berlin, Germany; Berlin-Brandenburg Center for Regenerative Therapies (BCRT), Charité - University Medicine Berlin, Germany; H.-D.V. and M.S.-H. contributed equally as senior authors

## Abstract

The field of gene therapy has been galvanized by the discovery of the highly efficient and adaptable site-specific nuclease system CRISPR/Cas9 from bacteria.^1,2^ Immunity against therapeutic gene vectors or gene-modifying cargo nullifies the effect of a possible curative treatment and may pose significant safety issues.^3-5^ Immunocompetent mice treated with CRISPR/Cas9-encoding vectors exhibit humoral and cellular immune responses against the Cas9 protein, that impact the efficacy of treatment and can cause tissue damage.^5,6^ Most applications aim to temporarily express the Cas9 nuclease in or deliver the protein directly into the target cell. Thus, a putative humoral antibody response may be negligible.^5^ However, intracellular protein degradation processes lead to peptide presentation of Cas9 fragments on the cellular surface of gene-edited cells that may be recognized by T cells. While a primary T cell response could be prevented or delayed, a pre-existing memory would have major impact. Here, we show the presence of a ubiquitous memory/effector T cell response directed towards the most popular Cas9 homolog from *Streptococcus pyogenes* (SpCas9) within healthy human subjects. We have characterized SpCas9-reactive memory/effector T cells (T_EFF_) within the CD4/CD8 compartments for multi-effector potency and lineage determination. Intriguingly, SpCas9-specific regulatory T cells (T_REG_) profoundly contribute to the pre-existing SpCas9-directed T cell immunity. The frequency of SpCas9-reactive T_REG_ cells inversely correlates with the magnitude of the respective T_EFF_ response. SpCas9-specific T_REG_ may be harnessed to ensure the success of SpCas9-mediated gene therapy by combating undesired T_EFF_ response *in vivo*. Furthermore, the equilibrium of Cas9-specific T_EFF_ and T_REG_ cells may have greater importance in *Streptococcus* pyogenes-associated diseases. Our results shed light on the T cell mediated immunity towards the much-praised gene scissor SpCas9 and offer a possible solution to overcome the problem of pre-existing immunity.

## Text

SpCas9 was the first Clustered Regularly Interspaced Short Palindromic Repeats (CRISPR) associated nuclease hijacked to introduce DNA double-strand breaks at specific DNA sequences.^1^ Through the ease of target adaption and the remarkable efficacy, it advanced to the most popular tool for re-writing genes in research and potential clinical applications. The major concern for clinical translation of CRISPR/Cas9 technology is the risk for off-target activity causing potentially harmful mutations or chromosomal aberrations.^2,7^ High-fidelity Cas9 enzymes were developed to reduce the probability of these events.^8^ Furthermore, novel Cas9-based fusion proteins allow base editing or specific epigenetic reprogramming without inducing breaks in the DNA.^9,10^ Most approaches are based on the original SpCas9 enzyme that originates in the facultatively pathogenic bacterium *Streptococcus (S.) pyogenes.* Every eighth school-aged child has an asymptomatic colonization of the faucial mucosa.^11^ S. pyogenes-associated pharyngitis and pyoderma are among the most common bacterial infection-related symptoms worldwide and can, sometimes lead to abysmal systemic complications.^12^ Due to the high prevalence of S. *pyogenes* infections, we hypothesized that SpCas9 could elicit an adaptive memory immune response in humans. Very recently, SpCas9-reactive antibodies but not SpCas9-reactive T cells were detected in human samples.^13^ The absence of detectable T cell reactivity in that study might be due to a sensitivity issue as only IFN-γ expression was analysed. Anti-SpCas9-antibodies should not impact the success of gene therapy, since usually SpCas9 is either protected by a vector particle or directly delivered into the targeted cells. In contrast, a pre-existing T cell immunity, particularly if tissue-migrating T_EFF_ cells are present, would result in a fast inflammatory and cytotoxic response to cells presenting Cas9 peptides on their major histocompatibility complexes (MHC)-molecules during or after intra-tissue gene editing.^4^

For detection of a putative SpCas9-directed T cell response, we stimulated human peripheral blood mononuclear cells (PBMCs) with recombinant SpCas9 and analysed the reactivity of CD3^+^4^+^/8^+^ T cells by flow cytometry with a set of markers for T cell activation (CD137, CD154) and effector cytokine production (IFN-γ, TNF-α, IL-2) (Fig. 1a, b, Extended Data Fig 1).^14,15^ We relied on protein uptake, processing and presentation of SpCas9 peptides by professional antigen-presenting cells (APCs) to both MHC I- and II within the PBMCs. Intriguingly, all donors evaluated showed specific memory/effector T cell activation upon SpCas9 stimulation indicated by CD137 (4-1BB) upregulation in both, CD4 and CD8, T cell compartments (Fig. 1a, b, d, e, Extended Data Fig. 1). After subtraction of background an average of 0.28% (range 0.03-1.02 %) and 0.44 % (range 0.6-1.3%) expressed CD137 within CD4^+^ and CD8^+^ T cells, respectively (Fig. 1e). By multiparameter analysis at single cell level, we detected Cas9-specific multi-potent T_EFF_ expressing at least one or even more effector cytokines (CD4^+^ > CD8^+^ T cells) (Fig 1 b, c, f). The expression of the lymph node homing receptor CCR7 and the leucocyte common antigen isoform CD45RO allows for dissection of the reactive T cell subsets (Extended Data Fig. 2a).^16^ Accordingly, we discovered that the majority of SpCas9-reactive T cells belongs to the effector-memory (CD4^+^ and CD8^+^) and terminally differentiated effector memory effector cells (T_EMRA_) (CD8^+^) pool implying repetitive previous exposure to SpCas9, comparable with memory T cell response to the frequently reactivated cytomegalovirus (CMV) (Extended Data Fig. 2b-e).^17^ The few cells within the naïve compartment might be related to stem cell memory T cell subset within this population.^18^

**Figure 1.**
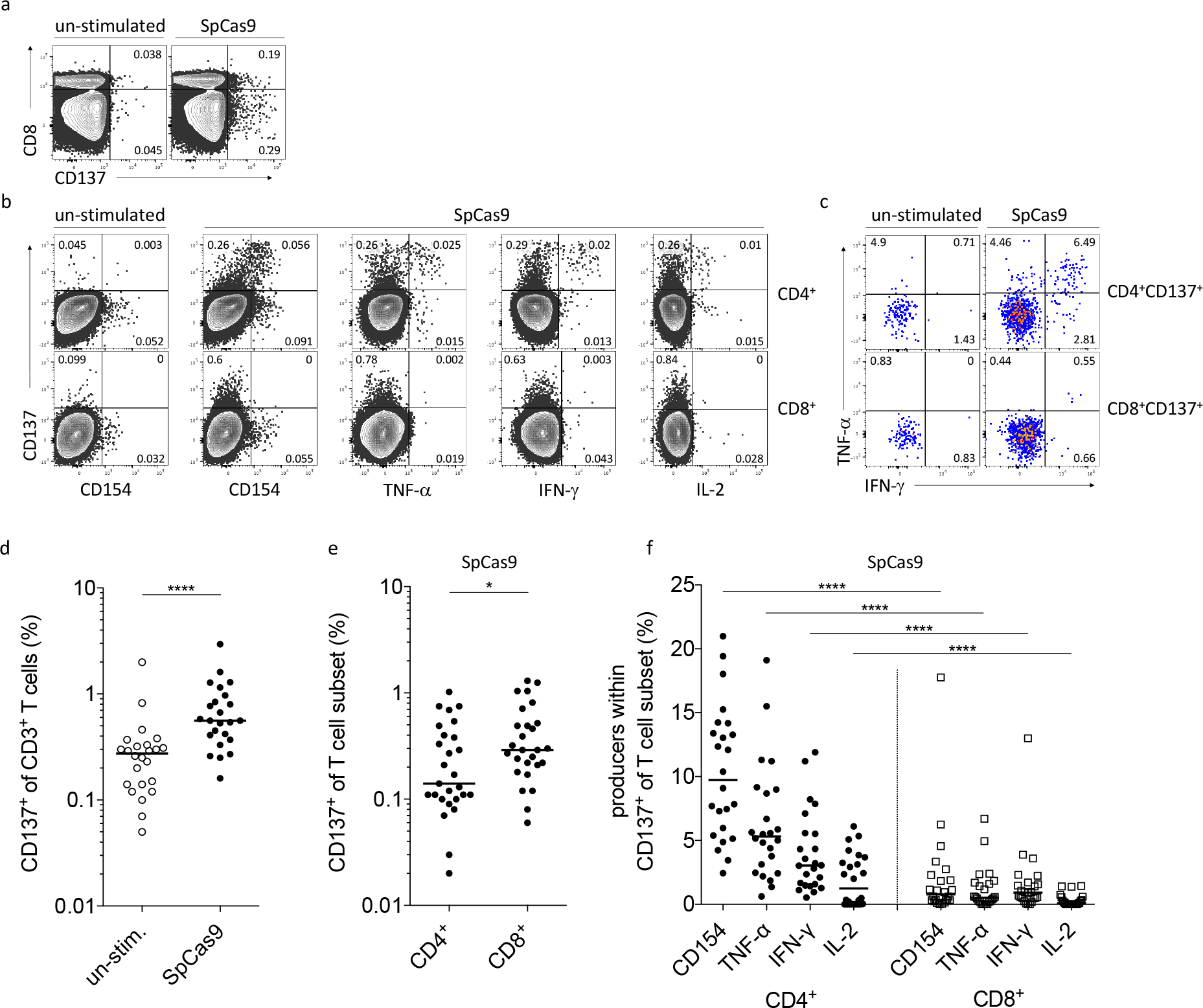
Ubiquitous peripheral SpCas9-specific T cell response within human donors. SpCas9-specific human CD3^+^ T cells can be identified after short-term *ex vivo* stimulation. PBMCs were stimulated with SpCas9 whole protein for 16 h. Frequencies of T cell response were assessed by flow cytometry, **(a)** Representative FACS plots show SpCas9-induced activation defined by CD137 expression of CD8^+^ and CD8^-^ T cells in comparison to unstimulated control. **(b)** Gating of single alive CD3^+^ T cells and dissection into CD4^+^ and CD8^+^ T cells. Representative FACS plots of SpCas9-induced CD137 and CD154 expression as well as IFN-γ, TNF-α and IL-2 production are shown. **(c)** Representative FACS plots of IFN-γ and TNF-α production within SpCas9-activated CD4^+^CD137^+^ and CD8^+^CD137^+^ T cells. **(d)** Paired analysis of SpCas9-induced CD137 expression within peripheral CD3^+^ T cells compared to unstimulated controls. **(e)** Background subtracted CD137 expression to SpCas9 whole protein by CD4+ and CD8+ T cells. **(f)** SpCas9-induced expression of CD154, TNF-α, IFN-γ and IL-2 within activated CD4^+^CD137^+^ and CD8^+^CD137^+^ T cells. (n=24; horizontal lines within graphs indicate medians.)

Our results imply a ubiquitous pre-primed T*EFF* response towards SpCas9, which could have immediate detrimental effects on tissues edited with a SpCas9-related system as those cells can immediately migrate to the targeted tissue. However, CMV is reactivated repeatedly in lymphoid organs and tissues, while S. *pyogenes* show repeated/continuous colonization on body surfaces. Recent studies indicate, that continuous colonialization and repetitive exposure to environmental proteins or pathogens particularly at mucosal surfaces also induce T_REG_.^19,20^ These T_REG_ are required to balance immune responses or even to maintain tolerance against innocuous environmental antigens.^20^ These findings expanded the significance of T_REG_ from controlling auto-reactivity towards a general role for protection against tissue-damaging inflammation. To determine the relative contribution of T_REG_to the SpCas9-induced T cell response, we performed intracellular staining for the T_REG_ lineage determining transcription factor FoxP3 in concert with CD25 surface expression.^21,22^ Further, we combined those T_REG_ defining markers with activation marker and cytokine profiling following SpCas9 whole protein stimulation (Fig. 2a, d, Extended Data Fig. 3). Intriguingly, we found excessive frequencies of T_REG_ within SpCas9-reactive CD4^+^CD137^+^ T cells ranging from 26.7-73.5% of total response (Fig. 2a, b). We confirmed T_REG_ identity through additional phenotypic marker combinations like FoxP^+^CTLA-4^+^ or CD127^low^CD25^high^ (Fig. 2a, Extended Data Fig. 3a, b) and epigenetic analysis of the T_REG_-specific demethylation region (TSDR demethylation: T_REG_ 83.7%; T_EFF_ 1.87%; n=1).^23,24^ Further investigation of the SpCas9-induced T cell activation revealed distinct T cell lineage determining transcription factor profiles. CD4^+^FoxP3^+^ T_REG_ were exclusively found within the CD137^dim^CD154^-^ population, while CD4^+^Tbet^+^ T_EFF_ comprised both CD137^+^CD154^+^ and CD137^high^ SpCas9-responsive populations (Fig. 2c, Extended Data Fig. 4). Functionally, T_REG_ did not contribute to SpCas9-induced effector cytokine production (Fig. 2d-f, Extended Data Fig. 5) but displayed a memory phenotype (Extended Data Fig. 3d). Taken together, our findings demonstrate that SpCas9-specific T_REG_ are an inherent part of the physiological human SpCas9-specific T cell response.

**Figure 2.**
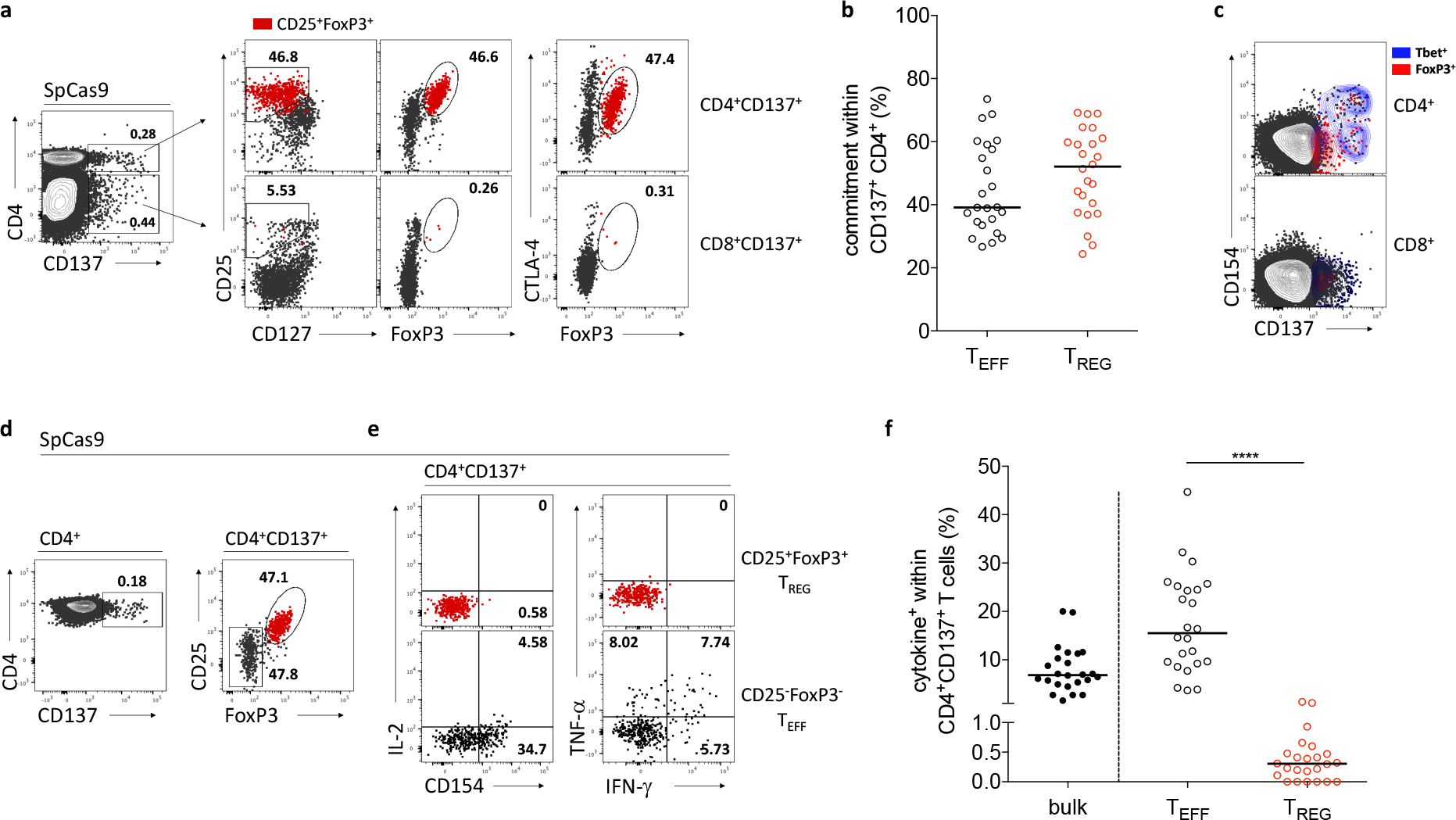
SpCas9-specific T cell response contains a substantial proportion of regulatory T cells. Identification of T_EFF_ and T_REG_ phenotypes within CD137^+^ T cells after 16 h stimulation of human PBMCs with SpCas9 whole protein. **(a)** Representative FACS plots show FoxP3 expression of T_REG_-defining markers CD25, FoxP3, CTLA-4 and CD127 within SpCas9-activated CD4^+^CD137^+^ and CD4^-^CD137^+^ T cells. The overlay highlighted in red represents CD25^+^FoxP3^+^ of CD137^+^ T cells. **(b)** Contribution to SpCas9-induced CD4^+^CD137^+^ T cell response by T_EFF_ and CD25^+^Foxp3^+^ T_REG_ phenotypes. **(c)** Overlay contour plots of a representative donor demonstrate Tbet^+^ (blue) and FoxP3^+^ (red) T cells within SpCas9-induced T cell activation defined by CD137 and CD154 expression. **(d)** Gating of CD4^+^T_REG_ within SpCas9-induced CD4^+^CD137^+^ T cells and **(e)** corresponding CD154 expression and cytokine production within CD4^+^CD137^+^ T_REG_ (red) and T_EFF_ (black). **(f)** Summary of accumulated cytokine production within bulk CD4^+^CD137^+^ T cells, CD4^+^CD137^+^ T_EFF_ (CD25^-^ FoxP3^-^) and CD4^+^CD137^+^ T_REG_ (CD25^+^FoxP3^+^). (n=24; horizontal lines within graphs indicate median values.)

Next, we investigated the individual relationship of T_EFF_ and their T_REG_ counterpart within the SpCas9-T cell response in comparison to an antiviral CMV and bacterial superantigen by relating the frequency of SpCas9, CMV phosphoprotein 65 (CMV_pp65_) and *Staphylococcus* Enterotoxin B (SEB)-activated T_REG_ to those of T_EFF_ within CD4^+^CD137^+^ and T_EFF_ within CD8^+^CD137^+^ antigen-reactive T cells. Remarkably, we found a balanced effector/regulatory T cell response to SpCas9 for both, CD4^+^ and CD8^+^, T cell compartments while response to CMV_pp65_ as well as SEB was dominated by T_EFF_ (Fig. 3a, b). Intriguingly, frequency of SpCas9-reactive CD4^+^CD137^+^CD154^-^ T_REG_ cells inversely correlates with the magnitude of CD4^+^CD137^+^CD154^+^ T_EFF_ within the SpCas9-reactive CD4^+^CD137^+^ T cells (Fig. 3c). In other words, our data show that donors with low SpCas9-reactive T_REG_ have relatively higher T_EFF_ response to SpCas9 suggesting that the level of SpCas9-specific T_EFF_ response might be controlled by SpCas9-specific T_REG_. A misbalanced SpCas9-reactive T_REG_/T_EFF_ ratio may result in an overwhelming effector immune response to SpCas9 following *in vivo* CRISPR/Cas9 gene editing.

**Figure 3.**
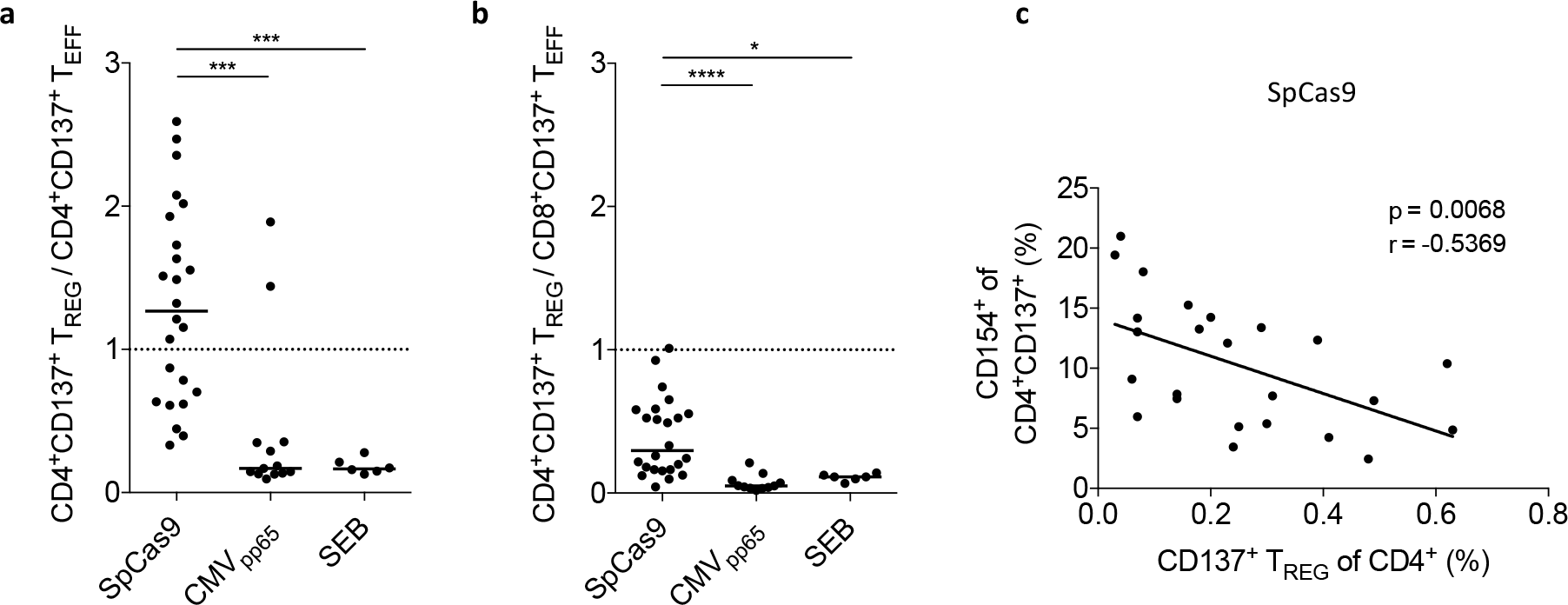
A balanced effector/regulatory T cell response to SpCas9 whole protein. (**a**) Relation of antigen-reactive T_REG_to CD4^+^T*EFF* shown for SpCas9 whole protein, CMV_*pp65*_ peptides and SEB stimulation. Antigen-reactive T_REG_ and T_EFF_ were defined according to gating strategy presented in Fig. 2d. Ratio was calculated by dividing the frequency of T_REG_ by the proportion of T_REG_ within CD4*+*CD137^+^ antigen-reactive cells. (**b**) Relation of antigen-reactive T_REG_ to CD8^+^T_EFF_ shown for SpCas9 whole protein, CMV_*pp65*_ peptides and SEB stimulation. Ratio was calculated by dividing the frequency of T_REG_ by the proportion of T_EFF_ within CD4^+^CD137^+^ antigen-reactive cells. (**c**) Inverse correlation of SpCas9-reactive T*REG* and SpCas9-reactive CD4^+^CD137^+^CD154^+^ T_EFF_. Pearson correlation coefficients were computed between frequency of SpCas9-reactive CD4^+^CD137^+^ T_REG_ within total CD4^+^ and the proportion of CD154^+^ cells within the SpCas9-activated CD4^+^CD137^+^ T cell pool. (SpCas9: n=24, CMV_pp65_: n=12, SEB: n=6. Horizontal lines within graphs indicate median values.)

Several preclinical and first clinical data show that adoptively transferred T_REG_ are able to combat not only T cell priming but also overwhelming T_EFF_ response.^25,26^Therefore, SpCas9-specific T_REG_ may have the potential to mitigate a SpCas9-directed T_EFF_ response. Having demonstrated that some individuals have a relatively low SpCas9-specific T_REG_/T_EFF_ ratio, adoptive transfer of those cells would be an option. Therefore, we tested enrichment and *in vitro* expansion of both SpCas9-specific T_EFF_ and T_REG_ (Extended Data Fig. 6). To examine their SpCas9-specific effector function, we re-stimulated T_EFF_ lines with SpCas9-loaded APCs after expansion and detected pronounced effector cytokine production (Extended Data Fig. 7). Notably, most cells within the SpCas9-specific T_REG_ lines lost their T_REG_-specific phenotype when cultured with IL-2, but were stabilized in the presence of the mTOR-inhibitor rapamycin, which is commonly used for expansion of thymic-derived naturally occurring T_REG_.^27^

What might be the physiological significance of a relatively high frequency of SpCas9-specific T_REG_ compared to CMV/SEB? Bacterial colonization requires homeostasis between the host and the microbiota for optimal coexistence. This interplay is tightly mediated by microbe-specific T_REG_. Prominently, patients suffering from immunodysregulation polyendocrinopathy enteropathy X-linked (IPEX) syndrome lacking functional T_REG_ cells fail to establish a healthy commensal flora resulting in multiple immunopathologies.^28^ Interestingly, *S. pyogenes* infection-associated diseases leading to systemic complications like rheumatic fever, occur predominantly in children and during adolescence.^12^ The pathophysiology is believed to involve molecular mimicry inducing cross-reactive antibodies by T helper cells (T_H_).^29^ However, T_H_mediated inflammation is controlled by T_REG_. Therefore, it would be worth to prove whether a misbalanced S. pyogenes-specific T_REG_/T_eff_response may be related to S. pyogenes-associated diseases.

In conclusion, our findings imply the requirement for controlling SpCas9 T_EFF_ response for successful CRISPR/Cas9 gene editing *in vivo.* It remains to be elucidated whether SpCas9-directed T cells can migrate into tissues relevant for gene therapy. Our results emphasize the necessity of stringent immune monitoring of SpCas9-specific T cell responses, preceding and accompanying clinical trials employing Cas9-derived therapeutic approaches to identify potentially high-risk patients. Henceforth, misbalanced T_REG_/T_EFF_ ratios and strong CD8^+^ T cell responses to SpCas9 may exclude patients for Cas9-associated gene-therapy. Gene editing with only transient SpCas9 exposure may reduce the risk for hazardous immunogenicity events. In contrast, technologies relying on *ex vivo* modification will not have a problem with immunogenicity because the gene-edited cells can be infused after complete degradation of the Cas9 protein. Unresponsiveness of autologous SpCas9-specific T_EFF_ lines to stimulation with CRISPR/Cas9-edited cell samples could be a release criterion for cell/tissue products in CRISPR/Cas9-related gene therapy (Extended Data Fig. 7). For *in vivo* application of CRISPR/Cas9, immunosuppressive treatment must be considered, especially if the control by T_REG_ is insufficient due to low T_REG_/T_EFF_ ratio. Immunosuppressive drugs discussed for AAV-related gene therapy in naïve recipients, such as CTLA4-IgG and low dose prednisone, are inadequate to control a pre-existing T_EFF_ response.^30^ Adoptive transfer of SpCas9-specific T_REG_ should be considered as an approach to prevent hazardous inflammatory damage to CRISPR/Cas9-edited tissues and would circumvent the need for global immunosuppression.

## Materials and Methods

### Cell preparation

We collected blood samples from healthy volunteers after obtaining informed consent. We separated PBMCs from heparinized whole blood from healthy donors at different days (median age: 30, range: 18-57, 12 female/ 12 male) by lymphoprep density gradient centrifugation with a Biocoll-separating solution. PBMCs were cultured in complete medium, comprising VLE-RPMI 1640 medium supplemented with stable glutamine, 100 U/ml penicillin, 0.1 mg/ml streptomycin (all from Biochrom, Berlin, Germany) and 10% heat-inactivated FCS (PAA).

### Flow cytometry analysis

We stimulated freshly isolated PBMCs in polystyrene round bottom tubes (Falcon, Corning) at 37 °C in humidified incubators and 5% CO_2_ for 16 h with the following antigens: 8 μg/ml Streptococcus pyogenes (Sp) CRISPR associated protein 9 (Cas9) (SpCas9) (PNA Bio Inc., CA, USA), 1 μSEB (Sigma) and CMV_pp65_ overlapping peptide pool at 1 μg/ml (15mer, 11 \ overlap, JPT Peptide Technologies, Berlin, Germany). For functional and phenotypic characterisation, 5x10^6^ PBMC / 1 ml complete medium were stimulated. For analysis of antigen-induced intracellular CD154 and CD137 expression and IFN-γ, TNF-α and IL-2 production, we added 2 μg/ml Brefeldin A (Sigma). To allow for sufficient SpCas9 antigenic APC processing and presentation, Brefeldin A was added for the last 10 h of stimulation. After harvesting, extracellular T cell memory phenotype staining was performed using fluorescently conjugated monoclonal antibodies for CCR7 (PE, clone: G043H7), CD45RA (PE-Dazzle 594, clone: HI100) and CD45RO (BV785, clone: UCHL1) for 30 min at 4 °C. In certain experiments CD25 (BD, APC, clone: 2A3), CD127 (Beckman Coulter, APC-Alexa Fluor 700, clone: R34.34) and CD152 (CTLA-4) (BD, PE-Cy5, clone: BNI3) antibodies were used to define T_REG_ specific surface molecule expression. To exclude dead cells, LIVE/DEAD Fixable Blue Dead Stain dye (Invitrogen) was added. Subsequently, cells were fixed and permeabilised with FoxP3/Transcription factor staining buffer set (eBioscience) according to the manufacturer’s instructions. After washing, we stained fixed cells for 30 min at 4 °C with the following monoclonal antibodies: FoxP3 (Alexa Fluor 488, clone: 259D), CD3 (BV650, clone: OKT3), CD4 (PerCp-Cy5.5, clone: SK3) CD8 (BV570, clone: RPA-T8), CD137 (PE-Cy7, clone: 4B4-4), CD154 (BV711, clone 24-31), IFN-γ (BV605, clone 4S.B3), TNF-α (Alexa Fluor 700, clone: MAb11) and IL-2 (BV421, clone MQ1-17H12)). In particular experiments, antibodies for intracellular fluorescence staining of Tbet (Alexa Fluor 647, clone: 4B10) and FoxP3 were used to define T cell lineage determining transcription factor expression levels. All antibodies were purchased from Biolegend, unless indicated otherwise. Cells were analysed on a LSR-II Fortessa flow cytometer (BD Biosciences) and FlowJo Version 10 software (Tree Star). For *ex vivo* analysis, at least 1x10^6^ events were recorded. Lymphocytes were gated based on the FSC versus SSC profile and subsequently gated on FSC (height) versus FSC to exclude doublets. Unstimulated PBMC were used as controls and respective background responses have been subtracted from SpCas9 or CMV_pp65_-specific cytokine production (Fig. 1d). Negative values were set to zero.

### SpCas9-specific T cell isolation and expansion

#### Isolation

We separated PBMCs from 80 mL heparinized whole blood. We washed PBMCs twice with PBS and cultured them for 16 h at 37 °C in humidified incubators and 5% CO_2_ in the presence of 8 μg/ml SpCas9 whole protein and 1 μg/ml CD40-specific antibody (Miltenyi Biotech, HB 14) at cell concentrations of 1x10^7^ PBMCs per 2 mL VLE-RPMI 1640 medium with stable glutamine supplemented with 100 U/ml penicillin, 0.1 mg/ml streptomycin and 5% heat-inactivated human AB serum (PAA) in polystyrene flat bottom 24 well plates (Falcon, Corning). After stimulation, cells were washed with PBS (0.5% BSA) and stained for 10 minutes with BV650-conjugated CD3-specific antibody, PerCp-Cy5.5-conjugated CD4-specific antibody, APC-conjugated CD25-specific antibody, APC-Alexa Fluor 700-conjugated CD127-specific antibody (Beckman Coulter), PE-Cy7-conjugated CD137-specific antibody and BV711-conjugated CD154-specific antibody. SpCas9-specific T_REG_ (Extended Data Fig. 6a: CD3^+^CD4^+^CD137^+^CD154^-^CD25^high^CD127^-^) and SpCas9-specific T_EFF_ (Extended Data Fig. 6a: CD3^+^CD137^+^CD154^+^CD25^low^) were enriched by fluorescently activated cell sorting on a BD FACSAriaII SORP (BD Biosciences). In addition, polyclonal (pc) T_REG_ (Extended Data Fig. 6a: CD3^+^CD4^+^CD137^-^CD154^-^CD25^high^CD127^-^) and pc T_EFF_ (Extended Data Fig. 6a: CD3^+^CD137^+^CD154^+^CD25^low^) were enriched for non-specific expansion. Intracellular T_REG_-specific FoxP3 transcription factor staining was performed post-sorting. Post-sorting analysis of purified subsets revealed greater than 90% purity.

#### Expansion

We cultured isolated SpCas9-specific T_EFF_ and control pc T_EFF_ cells at 37 °C in humidified incubators and 5% CO_2_ at a ratio of 1:50 with irradiated autologous PBMC (30 gy) in a 96-well plate (Falcon, Corning) with RPMI medium containing 5% human AB serum including 50 U/mL recombinant human (rh) IL-2 (Proleukin, Novartis). Isolated SpCas9-specific T_REG_ cells were cultured at 37 °C in humidified incubators and 5% CO_2_ at a ratio of 1:50 with irradiated autologous PBMC (30 gy) in a 96-well plate with X-Vivo 15 Medium (Lonza) containing 5% human AB serum including 500 U/mL rh IL-2 in the presence or absence of 100nM rapamycin (Pfizer). Non-specific pc T_REG_ were activated for polyclonal expansion applying the T_REG_ expansion kit according to the manufacturer’s instructions (T_REG_ : bead ratio of 1:1; CD3/CD28 MACSiBead particles, Miltenyi Biotech, Germany) and cultured in X-Vivo 15 Medium in the presence of 100nM rapamycin. We isolated a minimum of 10^4^ SpCas9-specific CD137^+^CD154^-^ T_REG_ cells, which could be expanded in vitro to at least 10^5^ cells within 10 days. Medium and cytokines were added every other day or when cells were split during expansion.

### *In vitro* restimulation of *ex vivo* isolated and expanded SpCas9-specific T cells

Cultured SpCas9-specific T_EFF_ and T_REG_ were analysed at day 10 for expression of effector molecules in response to stimulation with SpCas9 whole protein-loaded autologous monocyte-derived dendritic cells (moDCs). CD14^+^ monocytes were enriched from PBMCs by magnetically activated cell sorting (MACS, Miltenyi Biotech). Subsequently, CD14^+^ cells were cultured for 5 days in 1,000IU/mL rhGM-CSF (Cellgenix) and 400IU/mL rhIL-4 (Cellgenix). Then, fresh medium with 1,000IU/ml TNF-α (Cellgenix) was supplied. During 48 h of TNF-α induced maturation of autologous 4 μg/ml SpCas9 was added. We re-stimulated expanded T cell subsets with either SpCas9-pulsed, 1 μg/ml CMV_pp65_ overlapping peptide pool-pulsed or un-pulsed autologous moDCs for 6 h at a ratio of 10:1. 2 μg/ml Brefeldin A was added for the last 5 h of stimulation. Following stimulation, we analysed the expression of CD3, CD4, CD8, CD25, intracellular IFN-γ, TNF-α and IL-2, and intra-nuclear FoxP3, and treated the cells for flow cytometric readout as described above. We stained cells with BV650-conjugated CD3-specific antibody, PerCp-Cy5.5-conjugated CD4-specific antibody, BV570-conjugated CD8-specific antibody, APC-conjugated CD25-specific antibody, BV605 conjugated IFN-γ-specific antibody, Alexa Fluor 700 conjugated TNF-α-specific antibody and BV421-conjugated IL-2-specific antibody.

### TSDR - Methylation analysis

DNA methylation analysis of the T_REG_-specific demethylation region (TSDR) was performed as previously described.^24^ Briefly, bisulfite-modified genomic DNA (Quick-DNA Miniprep Plus Kit, Zymo Research, Irvine, USA; EpiTect Bilsulfite Kit, Qiagen, Hilden, Germany) was used in a real-time polymerase chain reaction for FoxP3 TSDR quantification. A minimum of 40 ng genomic DNA or a respective amount of plasmid standard was used in addition to 10 μl FastStart Universal Probe Master (Roche Diagnostics, Mannheim, Germany), 50 ng/μl Lambda DNA (New England Biolabs, Frankfurt, Germany), 5 pmol/μl methylation or nonmethylation-specific probe, 30 pmol/μl methylation or nonmethylation-specific primers (both from Epiontis, Berlin, Germany) in 20 μl total reaction volume. The samples were analysed in triplicate on an ABI 7500 cycler (Life Technologies Ltd, Carlsbad, USA).

### Statistical analysis and calculations

Graph Pad Prism version 7 was used for generation of graphs and statistical analysis. To test for normal Gaussian distribution Kolmogorov-Smirnov test, D’Agostino & Pearson normality test and Shapiro-Wilk normality test were performed. In two data set comparisons, if data were normally distributed Student’s paired t test was employed for analysis. If data were not normally distributed Wilcoxon’s matched pairs test was applied. All tests were two-tailed. Where we compared more than two paired data sets, one way ANOVA was employed for normally distributed samples and Friedman’s test was used for not normally distributed samples. For comparison of more than two unpaired not normally distributed data sets, we applied Kruskal-Wallis’ test. To exactly identify significant differences in not normally distributed data sets Dunn’s multiple comparison test was used as post-test and the post-test employed for normally distributed samples was Tukey’s multiple comparison test. Correlation analysis was assessed by Pearson’s correlation coefficients for normally distributed data or non-parametric Spearman’s rank correlation for not normally distributed data. The regression line was inserted based on linear regression analysis. Probability (p) values of ≤0.05 were considered statistically significant and significance is denoted as follows: * = p ≤ 0.05; ** = p ≤ 0.01; *** = p ≤ 0.001; **** = p ≤ 0.0001.

## Acknowledgments

We would like to acknowledge the assistance of the BCRT Flow Cytometry Core Lab, Dr. D. Kunkel and J. Hartwig.

## Author contributions

D.L.W. led the project, designed the research, performed most of the experiments, analysed and interpreted the data, and wrote the manuscript. L.A. and D.J.W. established methods, performed some of the experiments and revised the manuscript. P.R. wrote the manuscript and supplied reagents. H.-D.V. designed the research, interpreted the data and wrote the manuscript. M.S.-H. led the project, designed the research, analysed and interpreted the data, and wrote the manuscript.

---

**Extended Figure 1.**
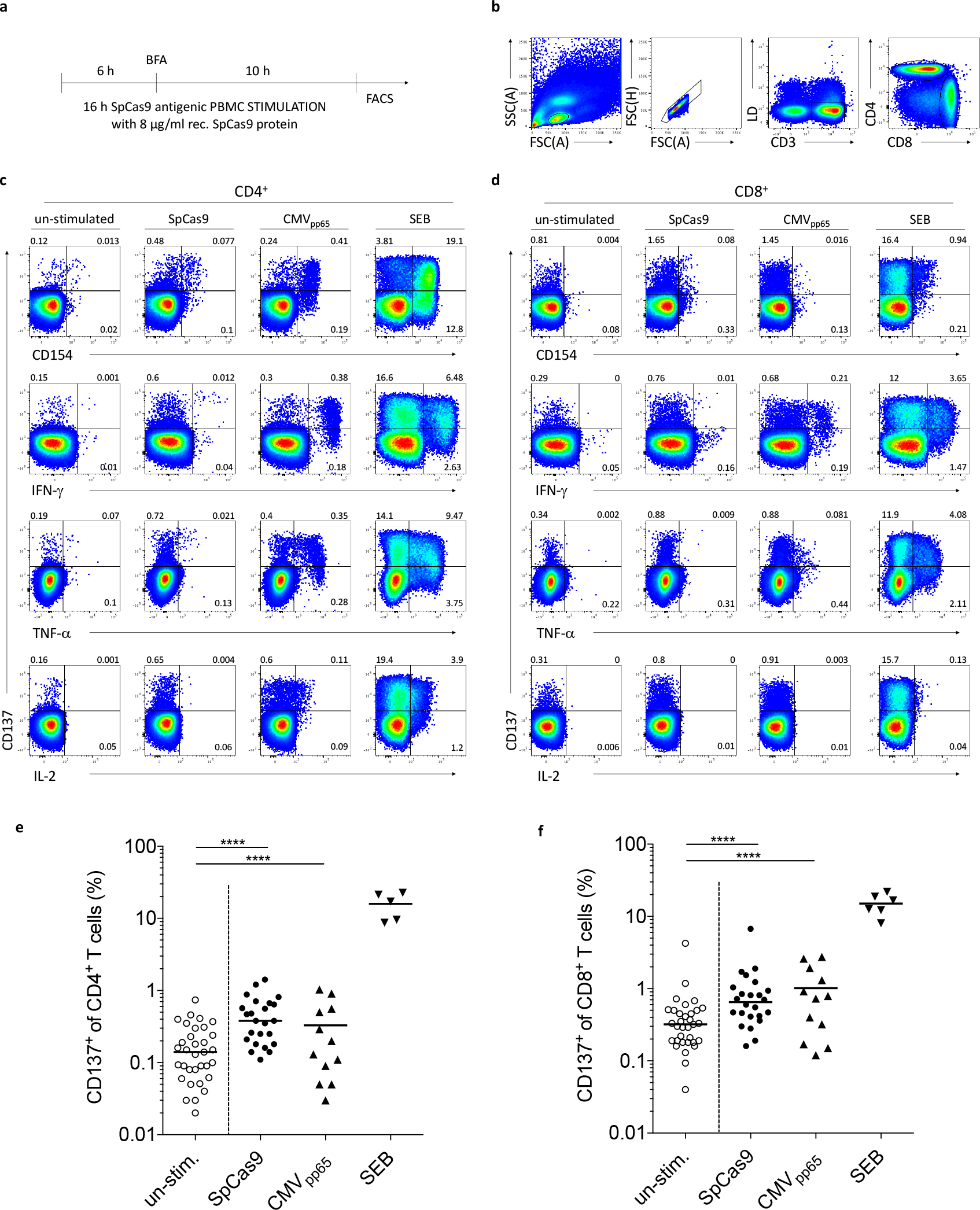
*Ex vivo* stimulation with SpCas9 whole protein induces polyfunctional effector CD4^+^ and CD8^+^ T cell responses. **(a)** Experimental design for ex *vivo* detection of SpCas9-specific *T* cell responses. **(b)** Representative gating strategy for defining alive CD3^+^CD4^+^ and CD3^+^CD8^+^ *T* lymphocytes. Lymphocytes were gated based on the FSC *versus* SSC profile and subsequently gated on FSC (height) *versus* FSC to exclude doublets. **(c** and **d;** summarized in **e** and **f)** Representative FACS images show SpCas9-induced activation defined by CD137 expression plotted against CD154, IFN-γ, TNF-α and IL-2 for CD4^+^ and CD8^+^ T cells in comparison to CMV_PP65_-stimulated and SEB-stimulated PBMCs. (SpCas9: n=24, CMV_PP65_: n=12, SEB: n=6. Horizontal lines within graphs indicate medians.)

**Extended Figure 2.**
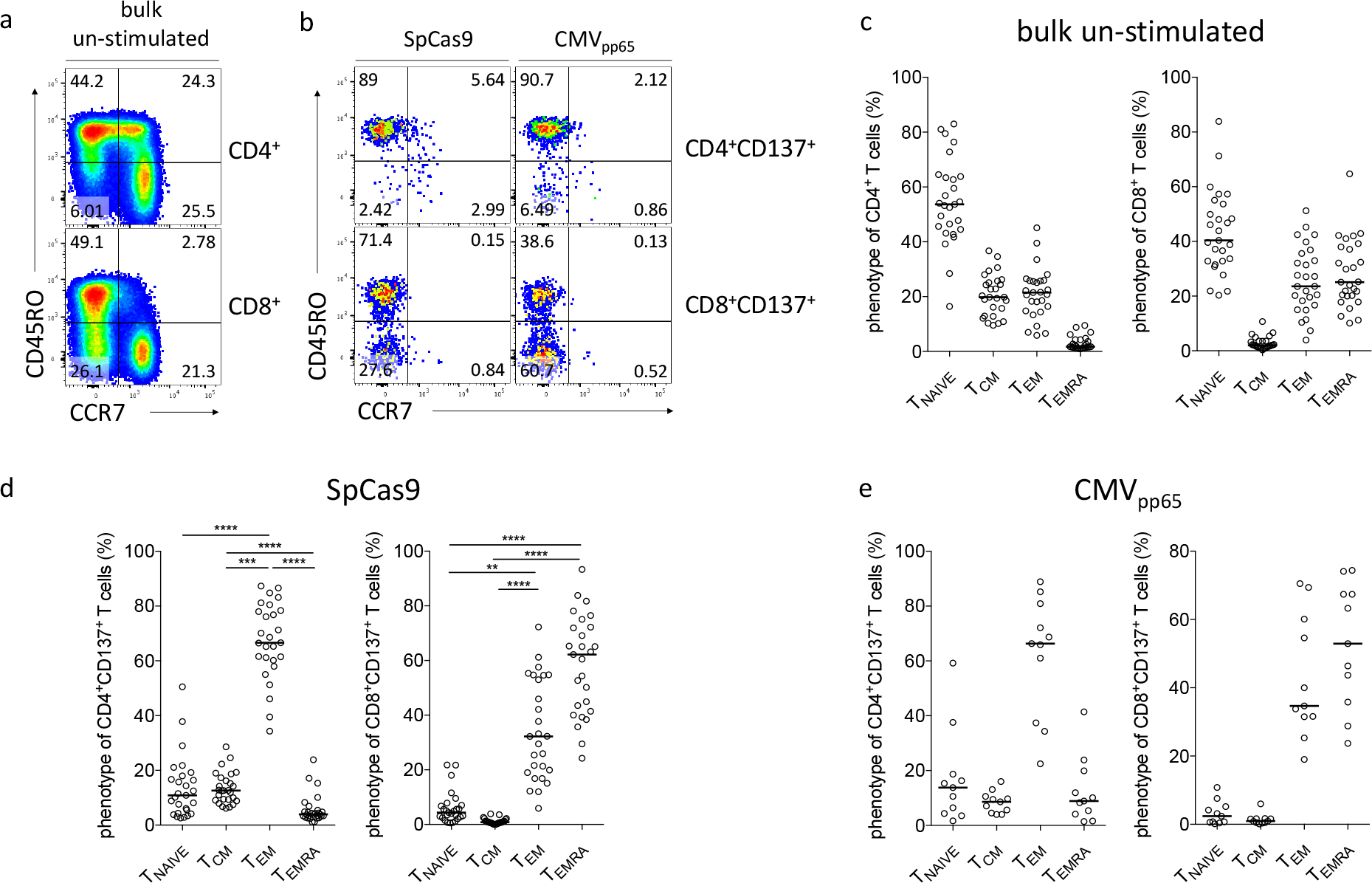
SpCas9- and viral CMV_PP_65-reactive CD4^+^ and CD8^+^ T cells phenotypically show a memory profile. (a) Strategy for defining T cell subsets from PBMCs according to the expression of CD3^+^ CD45RO^+^ and CCR7^+^ within CD4^+^ and CD8^+^ T cells. Dissection of the T cell differentiation profile into the following subsets: Naïve T cells (T_NAIVE_: CCR7^+^CD45RO^-^), central memory (T_CM_: CCR7^+^CD45RO^+^), effector memory (T_EM_: CCR7"CD45RO^+^) and terminally differentiated effector T cells (T_EMRA_: CCR7^-^CD45R^-^). **(b)** Strategy for defining T cell differentiation phenotypes applied to antigen-reactive CD4^+^CD137^+^ and CD8^+^CD137^+^ T cells after SpCas9 or human CMV_PP65_ PBMCs stimulation. Summarized phenotypical distribution of **(c)** bulk un-stimulated, **(d)** SpCas9-reactive (CD137^+^) and **(e)** CMV_PP65_-reactive (CD137^+^) CD4^+^ and CD8^+^ T cells. Flow cytometric analysis of PBMCs from a representative donor. (SpCas9: n=24. CMV_PP65_: n=10. Horizontal line in graphs indicates median value.)

**Extended Figure 3.**
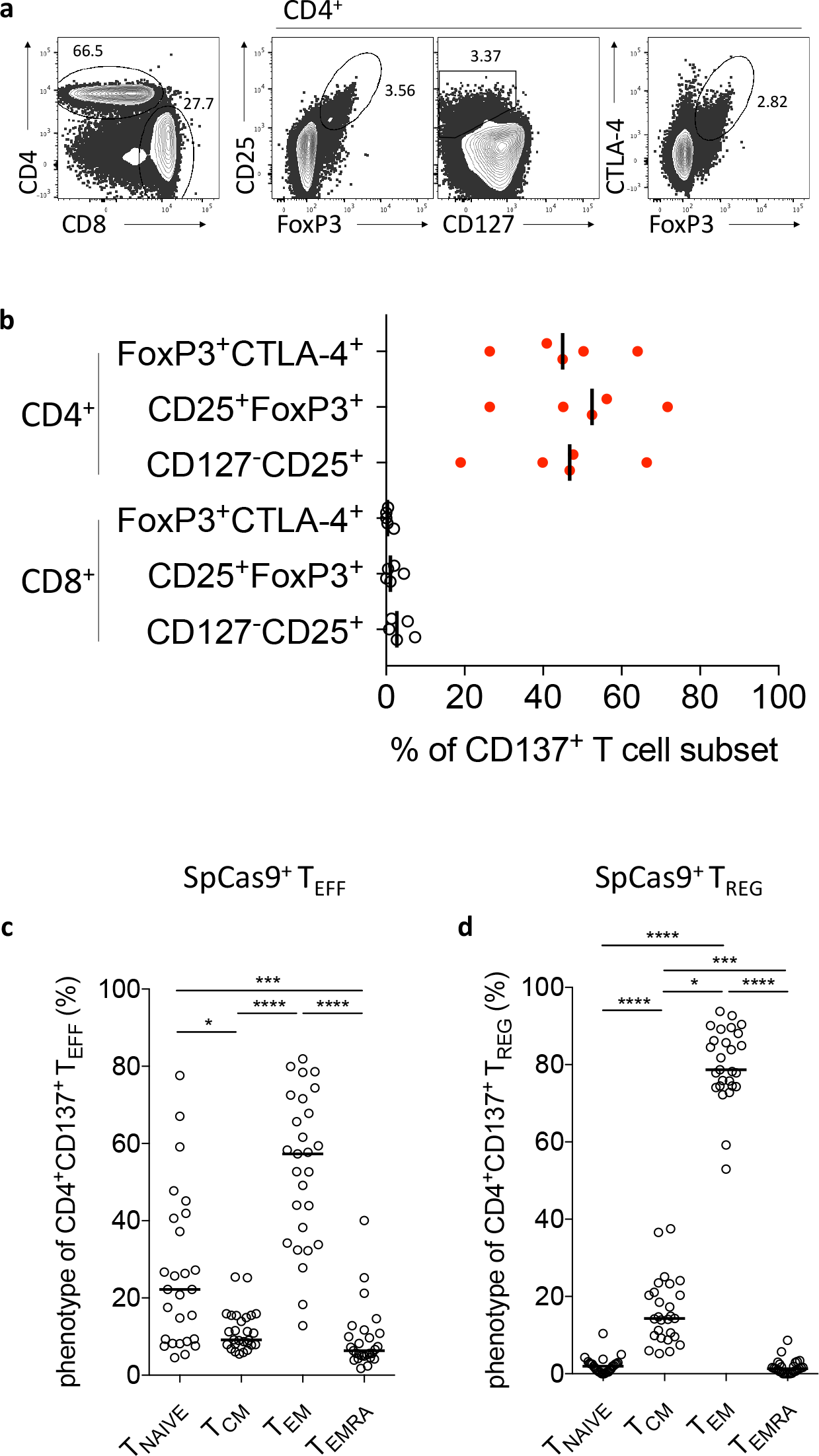
SpCas9-reactive CD4^+^CD137^+^ regulatory T cells show a memory phenotypic profile. **(a)** Gating strategy for the identification of T_REG_ phenotypes within the CD4^+^ T cell response. (b) Summary of T_REG_-defining markers CD25, FoxP3, CTLA-4 and CD127 within SpCas9-activated CD4^+^CD137^+^ and CD8^+^CD137^+^ T cells. **(c** and **d)** Summary of T cell differentiation phenotypes within SpCas9-reactive CD4^+^CD137^+^FoxP3^-^ T_EFF_ and CD25^+^Foxp3^+^ T_REG_. (n=24. Horizontal lines in graphs indicate median values.)

**Extended Figure 4.**
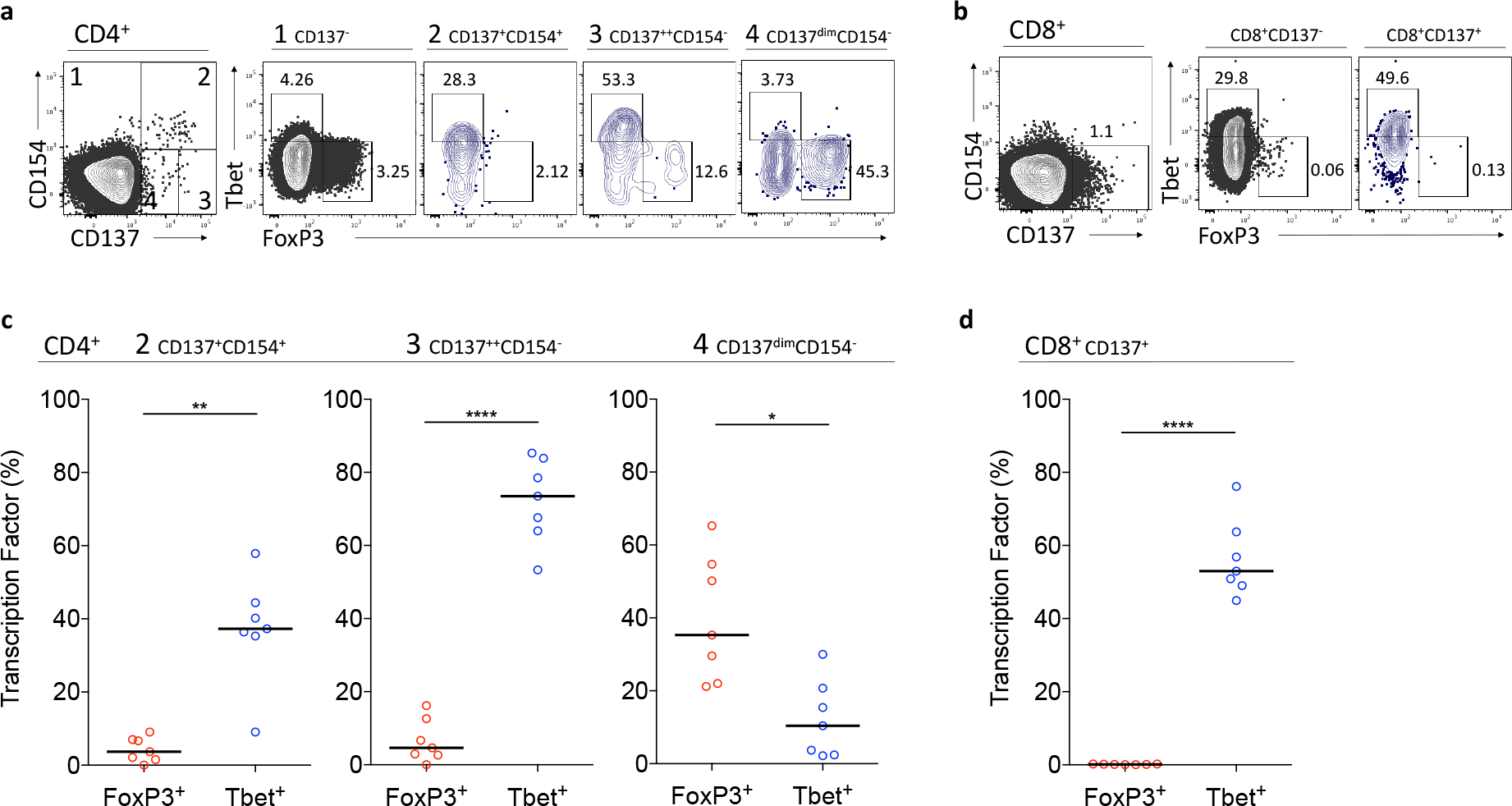
SpCas9-induced CD137 and CD154 expression correlate with lineage determining transcription factor pattern. The SpCas9-induced activation pattern on CD4^+^ was dissected according to CD137 and CD154 expression levels: **(1):** CD137^-^, **(2)** CD137^+^CD154^+^, **(3)** CD137^high^CD154^-^ and **(4)** CD137^dim^CD154^-^. SpCas9-reactive CD8^+^ T cells were defined through CD137 expression. Identification of Tbet (T_EFF_) and FoxP3 (T_REG_) transcription factors within **(a)** the CD4^+^ T cell response (1 to 4) and **(b)** the CD8^+^ T cell response to 16 h stimulation of human PBMCs with SpCas9 whole protein. **(c** and **d)** Summary of Tbet and FoxP3 expression within SpCas9-activated CD3^+^ T cells with designated activation pattern (CD4^+^: 2 to 4; CD8^+^: CD137^+^). (n=6; horizontal lines within graphs indicate median values.)

**Extended Figure 5.**
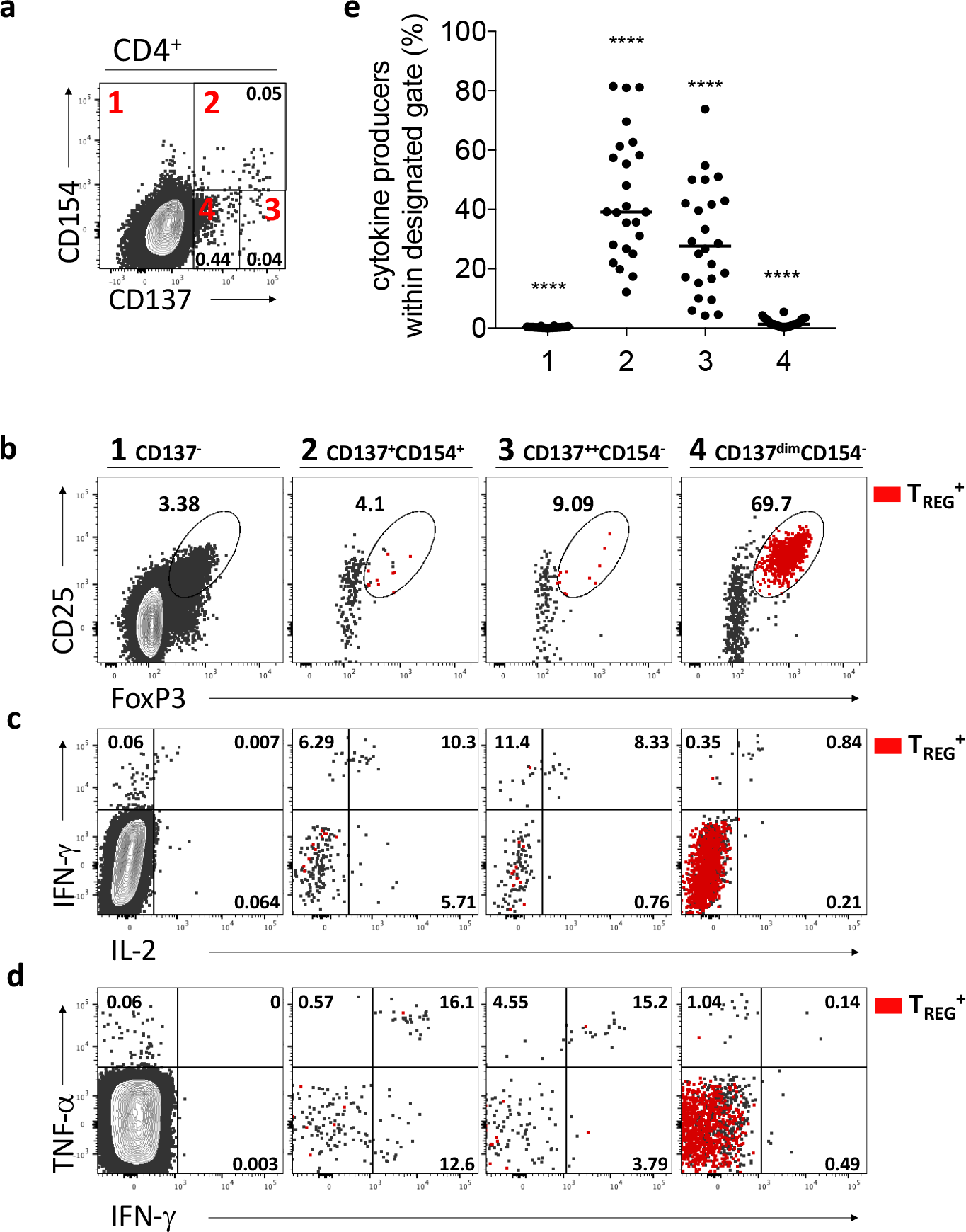
SpCas9-reactive CD4^+^ regulatory T cells are CD137^dim^ and lack CD154 expression and effector cytokine production. SpCas9-induced activation pattern on CD4^+^ T cells was dissected according to CD137 and CD154 expression levels: **(1):** CD137^-^, **(2)** CD137^+^CD154^+^, **(3)** CD137^high^CD154^-^ and **(4)** CD137^dim^CD154. **(a)** Representative FACS plots for SpCas9-induced activation pattern (1-4) and corresponding **(b)** T_REG_ phenotype (CD25^+^Foxp3^+^) and (**c** and **d**) effector cytokine production. Overlay demonstrates T_REG_ contribution to the SpCas9-induced T cell response (red). **(e)** Summary of accumulated cytokine production within T cells with designated activation pattern (1 to 4).

**Extended Figure 6.**
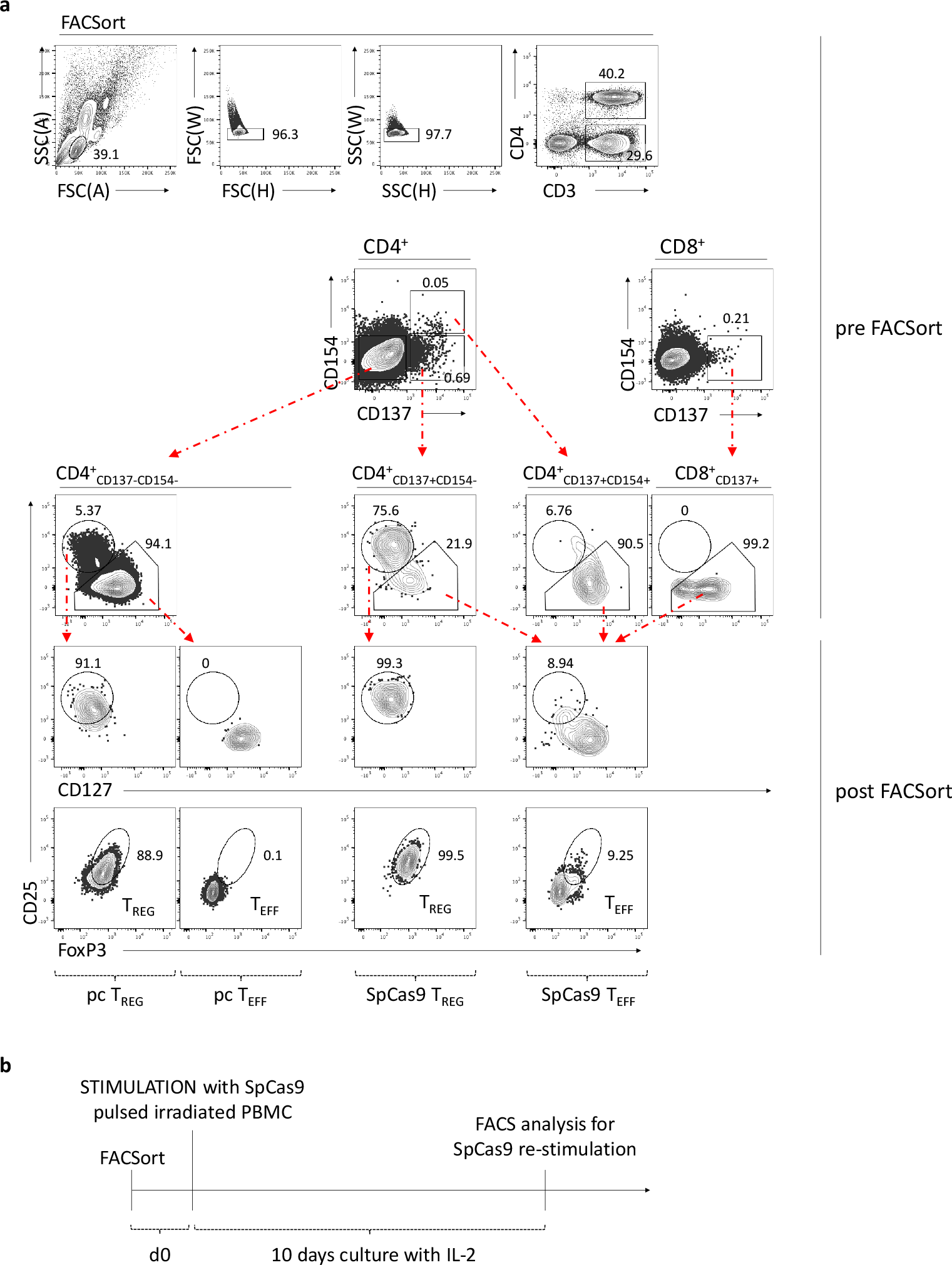
Flow cytometric enrichment of SpCas9-reactive T_EFF_and T_REG_. PBMCs were cultured for 16 h in the presence of 8 μg/ml SpCas9 whole protein and 1 μg/ml CD40-specific antibody. **(a)** SpCas9-specific T_REG_/T_EFF_ and un-stimulated pc T_REG_/T_EFF_ were enriched by FACSorting according to the incremental gating of CD3^+→^CD4^+^ or CD8^+→^ CD137^+/-^ CD154^+/-^ or CD137^+/-→^ CD25^high/low^CD127^+/-^. Post-sorting purity is shown in lower panels for CD4^+^CD137^-^CD154^-^CD25^high^CD127^-^ (pc T_REG_), CD4^+^CD137^-^CD154^-^CD25^loW^ and CD8^+^CD137^-^ CD154^-^CD25^loW^ (pc T_EFF_), CD4^+^CD137^+^CD154^-^CD25^high^CD127p^-^ (SpCas9 T_REG_) and CD4^+^CD137^+^CD154^+^CD25^low^, CD4^+^CD137^+^CD154^-^CD25^low^ and CD8^+^CD137^+^ (SpCas9 T_EFF_). Representative flow cytometric images shown. (n=2). **(b)** Experimental design for expansion and re-stimulation of enriched SpCas9-reactive T_EFF_ and SpCas9-reactive T_REG_and respective pc control populations.

**Extended Figure 7.**
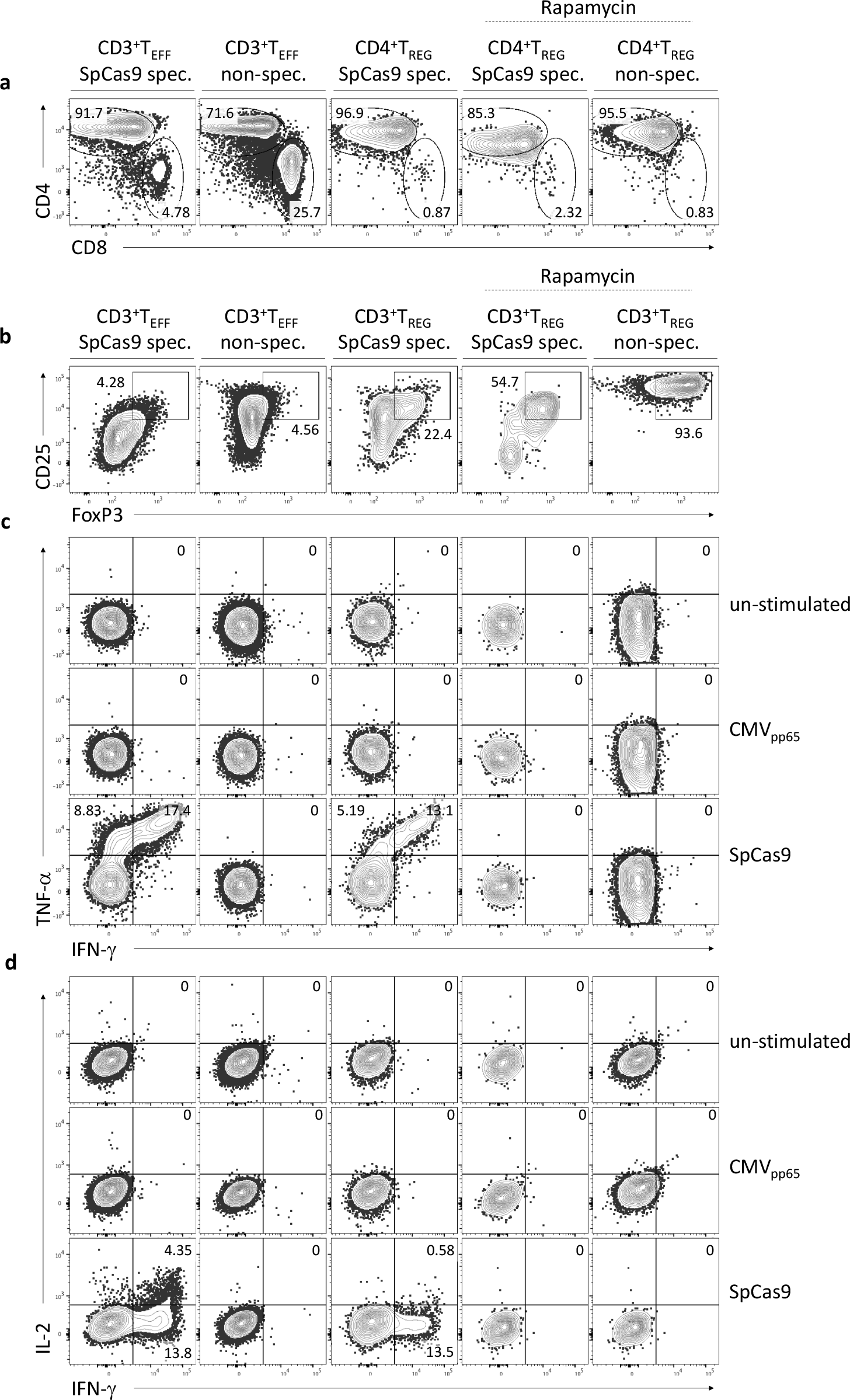
Expansion of SpCas9-reactive T cells. Antigen-specific readout for SpCas9-reactive *ex vivo* isolated and expanded T cells. Cultured SpCas9-specific T_EFF_ and T_REG_ were analysed at day *10* for expression of effector molecules in response to stimulation with SpCas9 whole protein loaded autologous moDCs for 6 h at a ratio of 10:1. Following stimulation, we analysed the expression of CD3, CD4, CD8, CD25, intracellular IFN-γ, TNF-α,IL-2 and FoxP3. **(a)** CD4 to CD8 ratio, **(b)** CD25 and FoxP3 expression, **(c)** TNF-_α_ and IFN-γ and **(d)** IFN-γ and IL-2 production within designated populations upon different stimuli (SpCas9, CMV_PP65_ and control).

